# Metabiosis underlies a microbiota permissive to Pseudomonadota and increases the risk of gut-borne bloodstream infection

**DOI:** 10.64898/2026.04.20.716137

**Authors:** Benjamin R. Wucher, Carlos J. Pardo-De la Hoz, Isaac Stamper, Sneh Sharma, Diego Kaune, Pranali Bendale, Jonathan Peled, Joao B. Xavier

## Abstract

The gut microbiota contains trillions of bacteria essential to health, but also harbors potential pathogens. The phylum Pseudomonadota, which includes *Escherichia coli*, *Klebsiella pneumoniae*, and *Pseudomonas aeruginosa*, typically composes <1% of the microbiota but causes disproportionate numbers of gut-borne bloodstream infections. Identifying the ecological dependencies that enable Pseudomonadota to cause gut-borne disease is important for human health. Here, we studied microbiota dynamics in patients undergoing allogeneic hematopoietic cell transplantation (allo-HCT) to find that microbiota compositions permissive to Pseudomonadota had, following antibiotic prophylaxis, high levels of *Bacteroides—*a major reservoir of polysaccharide utilization loci (PULs). We tested the causality of this clinical association in a mouse co-colonization model and discovered that *Bacteroides fragilis* promotes *Pseudomonas* gut colonization and survival to ciprofloxacin, a drug commonly used as prophylactic in allo-HCT. *In vitro* experiments revealed a general mechanism by which diverse Pseudomonadota species depend on *Bacteroides* polysaccharide breakdown to grow better, form more biofilm, and survive ciprofloxacin treatment under anaerobic conditions, a type of ecological dependency termed *metabiosis*. Guided by this insight, we used metagenomics to identify the PUL-encoded functions underlying the metabiotic potential of a patient’s microbiota and establish a link to gut-derived Gram-negative bacteremia in allo-HCT. Together, our findings translate mechanistically based microbiome ecology into a clinically actionable framework for early risk stratification and intervention.

## Introduction

The gut microbiota comprises trillions of bacteria from hundreds of diverse species. Most gut bacteria are beneficial to human health by aiding digestion^1–5^ and nutrient absorption^6–8^, or strengthening the immune system^6,9–11^, but some are potential pathogens. Gut-borne pathogens are typically present at low abundance and are kept from expanding by a healthy, biodiverse microbiota. However, treatments such as chemotherapy, which impair the immune system, and antibiotics, which disrupt the normal microbiota, can open ecological windows for pathogen expansion^12^. Identifying the processes that pathogenic bacteria can exploit is important for preventing gut-borne infections and informing treatment strategies, especially in immunocompromised people such as cancer patients receiving immunosuppressive therapy.

The phylum Pseudomonadota (formerly Proteobacteria) includes many genera and species with pathogenic potential, including Escherichia spp., Klebsiella spp., Pseudomonas spp., Enterobacter spp., Citrobacter spp., and Stenotrophomonas spp.^13,14^. These organisms are present in the microbiota of healthy people, but they typically constitute less than 1% of the bacterial population. Despite their low abundance, Pseudomonadota cause >50% of gut-borne bloodstream infections across patient cohorts^15^. This large contrast indicates that infection risk is not determined by the presence of Pseudomonadota alone, but by conditions that enable their expansion to a burden sufficient for gut-to-blood translocation^16^. These Gram-negative, facultative anaerobic pathogens are often natively resistant to many antibiotics, and increasing resistance to common treatments is occurring rapidly^13,17,18^.

Nonetheless, while antibiotic resistance may enable survival, it does not guarantee expansion in the gut, where the environment is anaerobic, and the dominant nutrients are complex dietary and host glycans that these organisms are poorly equipped to exploit^19^. Many of the carbon sources preferred by Pseudomonadota, like sugar monomers and amino acids are rapidly broken down by the host, making them unavailable in the colon.^20,21^

Under normal conditions, the gut microbiota is a biodiverse, functionally linked microbial ecosystem, in which many species compete intensely for nutrients^22^. This ecosystem is dominated by two phyla: the Bacteroidota (formerly Bacteroidetes) and the Bacillota (formerly Firmicutes). Primary utilizers, mostly members of Bacteroidota such as *Bacteroides*, can encode diverse polysaccharide utilization loci (PULs) that enable them to degrade complex dietary and host-derived glycans. This process releases sugars into the gut environment that can fuel secondary degraders, such as many commensal members of the Bacillota. This process is a form of metabiosis, in which one organism’s metabolic activity passively modifies the environment and generates the conditions needed for another to survive, without harm to either^7,23^. Pseudomonadota are also secondary degraders and compete with commensals for the metabiotic breakdown products of primary utilizers^24^. Ecosystem disturbances, such as antibiotics, could create ecological opportunities for Pseudomonadota by depleting consumer competitors while sparing key producers.

Allogeneic hematopoietic stem cell transplantation (allo-HCT) is a treatment for hematological malignancies that can cause immune deficiencies lasting days, weeks and sometimes months^25,26^. Concurrently, these patients also endure intestinal barrier damage during this period. As a result, allo-HCT patients are at high risk of bloodstream infections from the gut^16,27,28^. Fluoroquinolone prophylaxis (e.g., ciprofloxacin, levofloxacin), specifically administered to prevent Gram-negative infections, reduces the incidence of gut-borne bloodstream infections from 11% to 6%^15^, which remains a significant cause of transplant-related mortality^29^. A previous study of the microbiota dynamics in allo-HCT patients showed that intestinal expansion of Pseudomonadota increases the risk of Gram-negative bacteremia^15^. This clinical association suggests that routine longitudinal monitoring of fecal Pseudomonadota abundance could identify patients at high risk of bacteremia. However, the mechanisms that make the microbiome permissive to the expansion of Pseudomonadota in the first place remained unclear.

Here, we leveraged a large allo-HCT dataset^16^ to identify signatures of a permissive microbiota and found that high relative abundance of *Bacteroides* following antibiotic prophylaxis predispose patients to Pseudomonadota expansion and subsequent bloodstream infection. To assess the causality of this clinical finding, we investigated the interaction between *Pseudomonas* and *Bacteroides* in a mouse model and found that *Bacteroides* promotes *Pseudomonas* colonization and enhances its survival during ciprofloxacin treatment. Then, using a microfluidic device that mimics gut architecture, we found that *B. fragilis* enables protective, biofilm-like structures under anaerobic conditions. Complementary *in vitro* assays established a mechanism wherein different *Bacteroides* enhance the growth, biofilm formation, and ciprofloxacin tolerance of multiple Pseudomonadota species across a range of glycan substrates.

Importantly, this benefit was species-specific: different *Bacteroides* strains varied in their capacity to support Pseudomonadota depending on their PUL repertoires. Guided by this mechanism, we returned to the clinical cohort and identified commensal PUL functions that underlie the metabiotic potential driving microbiota permissiveness to Pseudomonadota and subsequent Gram-negative infection.

## Results

### Bacteroides post-prophylaxis is a signature of a microbiota permissive to Pseudomonadota

We analyzed a longitudinal microbiome dataset from patients undergoing allo-HCT (**Fig. 1A**), comprising 1318 transplant episodes (1276 patients) with fecal microbiome profiling in 1265 episodes across the day −15 to +35 peri-transplant window (example patient timeline in **Fig. 1B**). During this high-risk period across 1,265 evaluable transplantation episodes, there were 107 (8.1%) first Gram-negative bloodstream infection events in which a member of Pseudomonadota was identified in blood cultures (**Fig. 1C**; S1). Subsequent Gram-negative bloodstream infection in the same patients were excluded from the analysis.

**Fig. 1:**
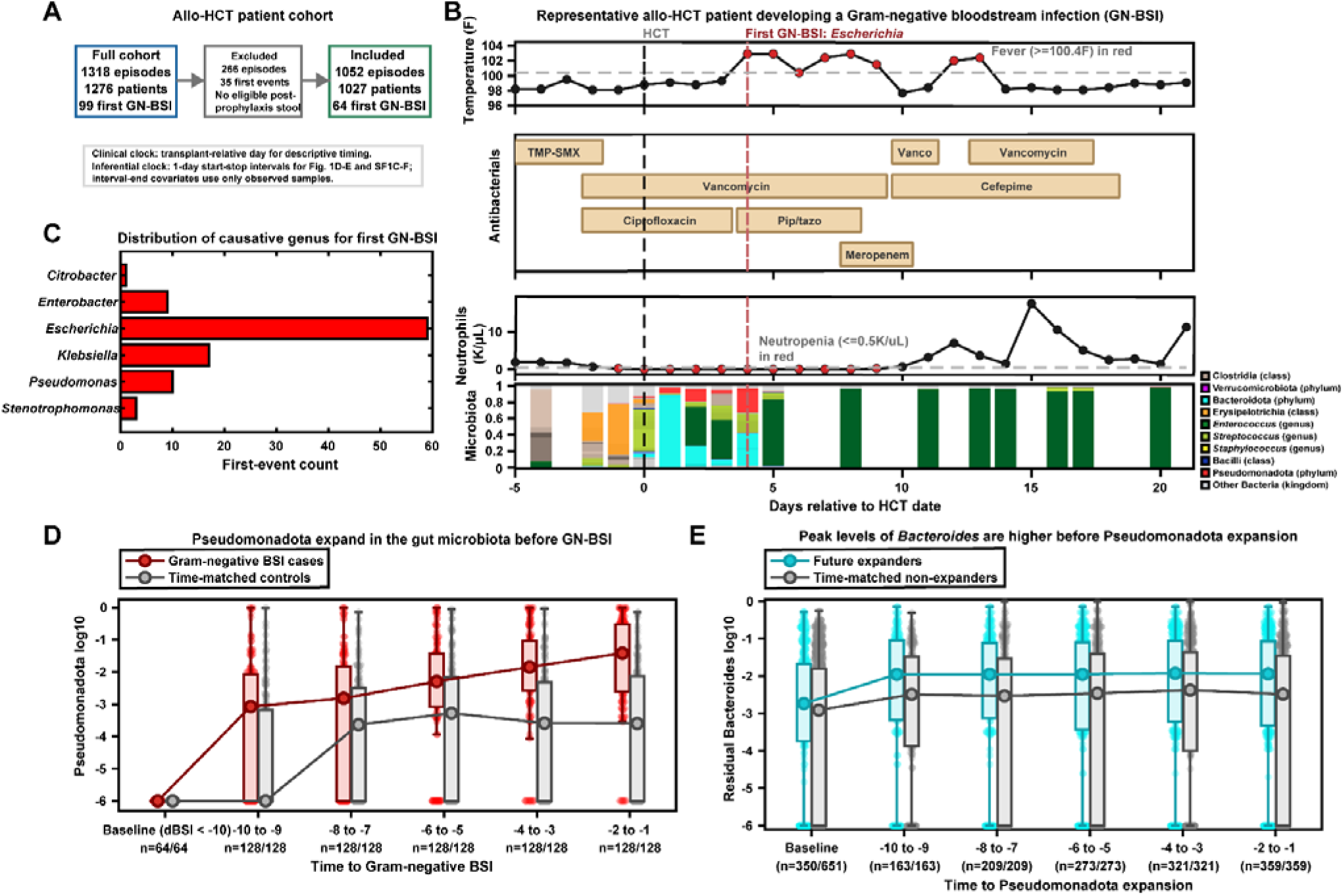
Post-prophylaxis Bacteroides Marks a Microbiota State Permissive to Pseudomonadota Expansion in Allo-HCT patients. (A) Cohort structure and leakage-safe inferential denominator. The cohort bridge defines the descriptive and inferential populations used in the section. The inferential denominator preserves the post-prophylaxis at-risk ecological window while excluding episodes that cannot support leak-free time-varying analysis. (B) Example patient timeline through the peri-transplant window. This single-patient timeline illustrates how fecal sampling, antibiotic exposure, blood counts, temperature, and bloodstream infection timing are aligned around transplant. The stool panel displays the top ASVs by overall abundance across the displayed stool samples. (C) Diverse causative genera underlie the first Gram-negative bloodstream infection. This supports a general ecological susceptibility to Pseudomonadota-linked gut-borne infection. (D) Fecal Pseudomonadota behaved as a proximate ecological state rather than a stable baseline discriminator. Baseline fecal Pseudomonadota burden was the same in future infection cases and controls, but in episodes that later developed bloodstream infection, fecal Pseudomonadota rose sharply as the event approached, while matched controls remained comparatively low. Late case samples were strongly shifted upward relative to baseline, while the case-control separation remained evident across the late pre-infection bins. Thus, the biologically relevant signal is not baseline carriage. It is a dynamic post-prophylaxis takeover state that emerges close to infection. (E) Residual *Bacteroides* diverges before expansion onset. Baseline uses pre-prophylaxis samples, and later bins show matched pre-expansion intervals. Jittered values are displayed over boxplots, and lines connect the within-bin medians. Future expanders already show higher residual *Bacteroides* than time-matched non-expanders before overt Pseudomonadota expansion.

Longitudinal sampling revealed a consistent dynamic preceding infection: among cases, Pseudomonadota increased as the infection approached, and sample-level distributions in the 15 days prior shifted upward relative to samples from non-infected episodes (**Fig. 1D**; 299 samples from 89 infection cases versus 7502 comparator samples from 1165 controls). Notably, baseline composition (microbiota composition before prophylaxis started) did not discriminate infected from non-infected cases: in 199/1318 episodes with baseline samples available (320 total), Pseudomonadota detection in baseline fecal samples was not associated with subsequent bloodstream infection after adjusting for baseline sequencing depth and sample count (OR ∼0.82–0.83; p>0.8). In contrast, the evolving Pseudomonadota abundance was a strong proximate risk marker: a time-varying Cox model showed a large increase in Gram-negative infection hazard with increasing fecal Pseudomonadota burden (HR ∼1.84 per +1 log10 unit). We therefore operationalized Pseudomonadota expansion as an intermediate ecological state (primary: first Pseudomonadota >5% after prophylaxis start). Expansion was detected in 383/1189 (32.2%) eligible episodes and was associated with a higher risk of bloodstream infection (10.7% with expansion vs 4.5% without; RR=2.40).

To identify specific signatures of a microbiota permissive to expansion, we analyzed the residual microbiota, with Pseudomonadota removed and the remainder renormalized, in expanding episodes versus time-matched non-expanding controls. In matched pre-expansion interval analyses, residual *Bacteroides* was a prominent component of this permissive microbiota, with higher abundances in future expanders than in time-matched non-expanders (**Fig. 1E**; OR = 1.16 per log_10_ unit, 95% CI 1.05-1.29, p = 5×10^-3^). This separation was already evident one week before expansion onset, when the median residual-*Bacteroides* signal differed by 0.5 log10 units. We then quantified permissiveness as an achieved state using the cumulative maximum of residual *Bacteroides* abundance after prophylaxis; this cummax residual-*Bacteroides* state was associated with increased hazard of subsequent Pseudomonadota expansion and remained directionally consistent across multiple expansion thresholds (for the primary >5% definition, HR = 1.28 per log_10_ unit, 95% CI 1.20-1.38, p = 4×10^-12^).

These results support a causal chain in which *Bacteroides* primarily contributes by increasing the hazard of entering the Pseudomonadota-expansion state; supporting the idea of a metabiotic pathway whereby post-prophylaxis *Bacteroides* increases gut permissiveness to expansion, raising the risk of bloodstream infection.

### Bacteroides supports Pseudomonas in antibiotic-treated mice

We then tested the causality of this clinical association using *Pseudomonas aeruginosa* as our model Pseudomonadota. This *Pseudomonas* species is normally absent from the microbiota of mice, but it causes gut-borne infections in allo-HCT^15,30^ with the highest mortality rate^30^. Because *P. aeruginosa* is known to secrete various toxins to eliminate competitors and to regulate their secretion in response to external stimuli^31^, we first ruled out the possibility that this model Pseudomonadota would simply kill the *Bacteroides*, which would invalidate our model choice. We treated *B. fragilis in vitro* with supernatants from *P. aeruginosa* cultured aerobically or anaerobically. The aerobically cultured supernatant prevented *B. fragilis* growth, while the anaerobic supernatant had no effect (**Fig. S2**, comparing *B. fragilis* growth in *Pseudomonas* supernatant from aerobic *vs* anaerobic culture). Therefore, the *Pseudomonas* antagonistic potential is context-dependent and is inactive under anaerobic conditions, supporting the notion that this species is a suitable model to study Pseudomonadota’s ability to exploit commensal populations.

We tested our ability to replicate the patient scenario, where the antibiotic-perturbed microbiota becomes permissive to *Pseudomonas,* which would then translocate and cause gut-borne infections. We first disturbed the native microflora of C57BL/6 mice with penicillin and streptomycin for 7 days, then gavaged a 1:1 mixture of *Bacteroides* and *Pseudomonas* (**Fig. 2A, cartoon**). Both cohorts show that Pseudomonas can colonize under these conditions as described previously^32^. However, compared with *Pseudomonas* alone, the co-inoculation with *Bacteroides* increased *Pseudomonas* load by a remarkable 100-fold (**Fig. 2B/D**). Subsequent ciprofloxacin treatment from day 15 to day 17 post-gavage (**Fig. 2A**, cartoon) had a strong and durable impact on the *Pseudomonas* load in mono-colonized mice. However, the ciprofloxacin had a much weaker effect in co-colonized mice, and the *Pseudomonas* levels recovered immediately after ciprofloxacin ceased (**Fig. 2B/D**). The *Bacteroides* load in both the monoculture and coculture with *Pseudomonas* remained stable for the entire experiment, again confirming that *Pseudomonas* does not harm *Bacteroides* within this context **(Fig. 2C).** After the experiment concluded, imaging of the intestines via fluorescence *in-situ* hybridization confirmed that both species could be found directly colocalizing within the colon **(Figure 2E)**.

**Figure 2:**
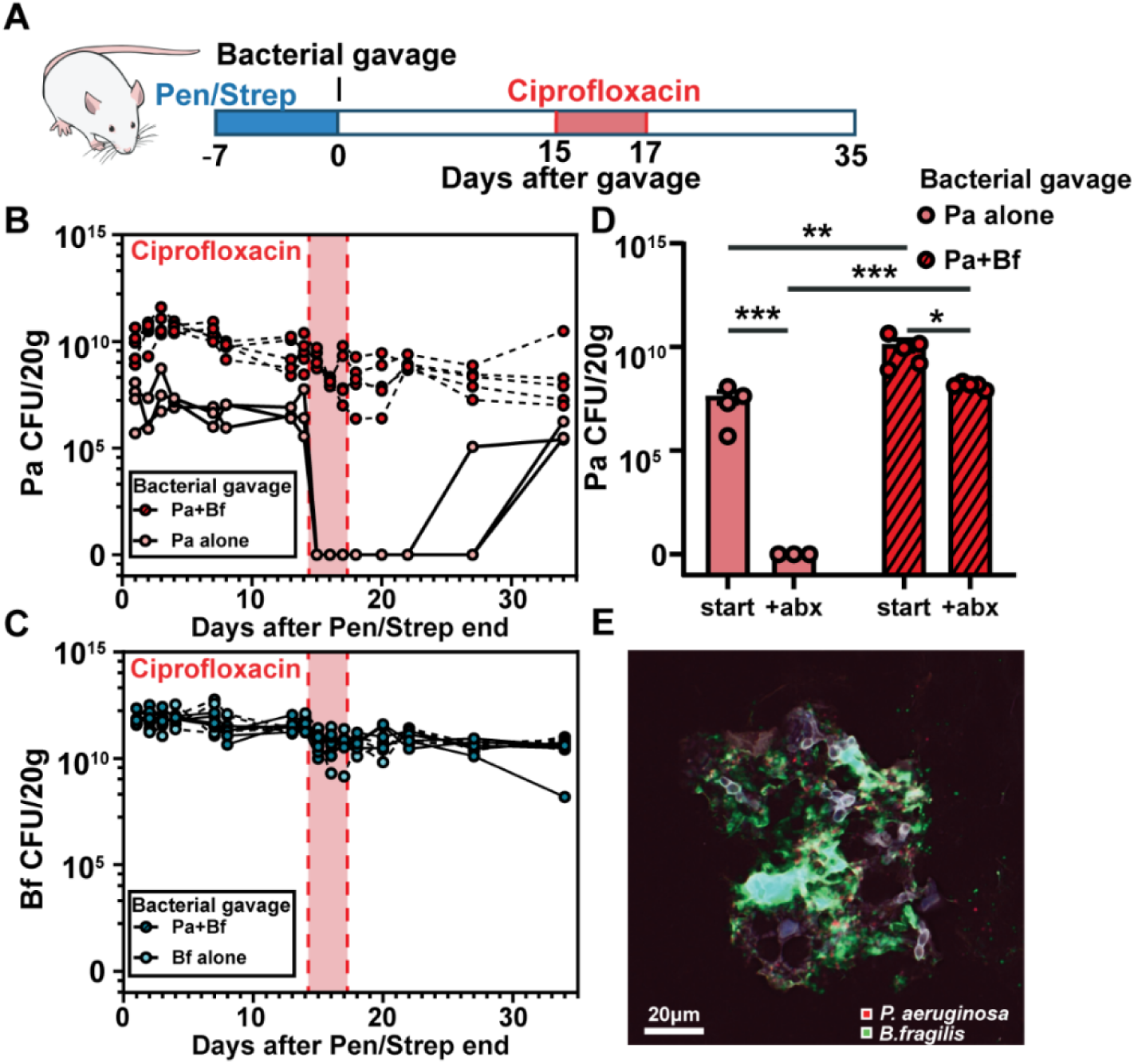
B*a*cteroides is highly present in the post antibiotic gut and can drastically impact pathogen colonization. (A) Outline of experimental protocol. Mice were first given a 7-day course of Penicillin and Streptomycin to reduce their gut microbial load. Complete schematic of the experiment timeline. The experiment mimics infection and treatment with the addition of a bacterial gavage and ciprofloxacin treatment monoculture n=4, coculture n=5. (B/C) A timeline showing *P. aeruginosa* and *B. fragilis* bacterial load in fecal pellets in both mono and coculture. Panels are split by species counts**. (D) Antibiotic effect on Pseudomonas.** Measurements of bacterial load at day 16 after gavage, Welch’s test *P<.05,**P<.01,***P<.001,****P<.0001. (E) *B. fragilis* and *P. aeruginosa* closely associate in the gut. Representative image from mouse colon at day 34 of *P. aeruginosa* and *B.fragilis* colocalization using FISH.

### Bacteroides boosts Pseudomonas biofilm and ciprofloxacin survival in anaerobic environments

Given the *in vivo* support for our patient findings, and the striking antibiotic tolerance of the co-culture, we next investigated the mechanism of interaction *in vitro*. We first used a microfluidic device to mimic the gut architecture by forming crypts with dimensions of approximately 80 μm wide and 400 μm long, in the range of the human intestine^33^ (**Fig. 3A**). We introduced mGFP-labelled *P. aeruginosa* into the device, either alone or in co-culture with mRFP-labelled *B. fragilis,* and incubated it for 48 hours under anaerobic conditions. For consistency of the color code used in our patient data, the *Pseudomonas* is pseudo-colored red (the color used to represent Pseudomonadota) and the *Bacteroides* is pseudo-colored cyan. We used a *Bacteroides*-specific minimal medium supplemented with 5mg/mL cyclodextrin, a complex polysaccharide made of glucose monomers joined together by α-1,4-glycosidic bonds in a cyclic structure, as the sole carbon source. Following incubation, the devices were removed and imaged via confocal microscopy.

**Figure 3.**
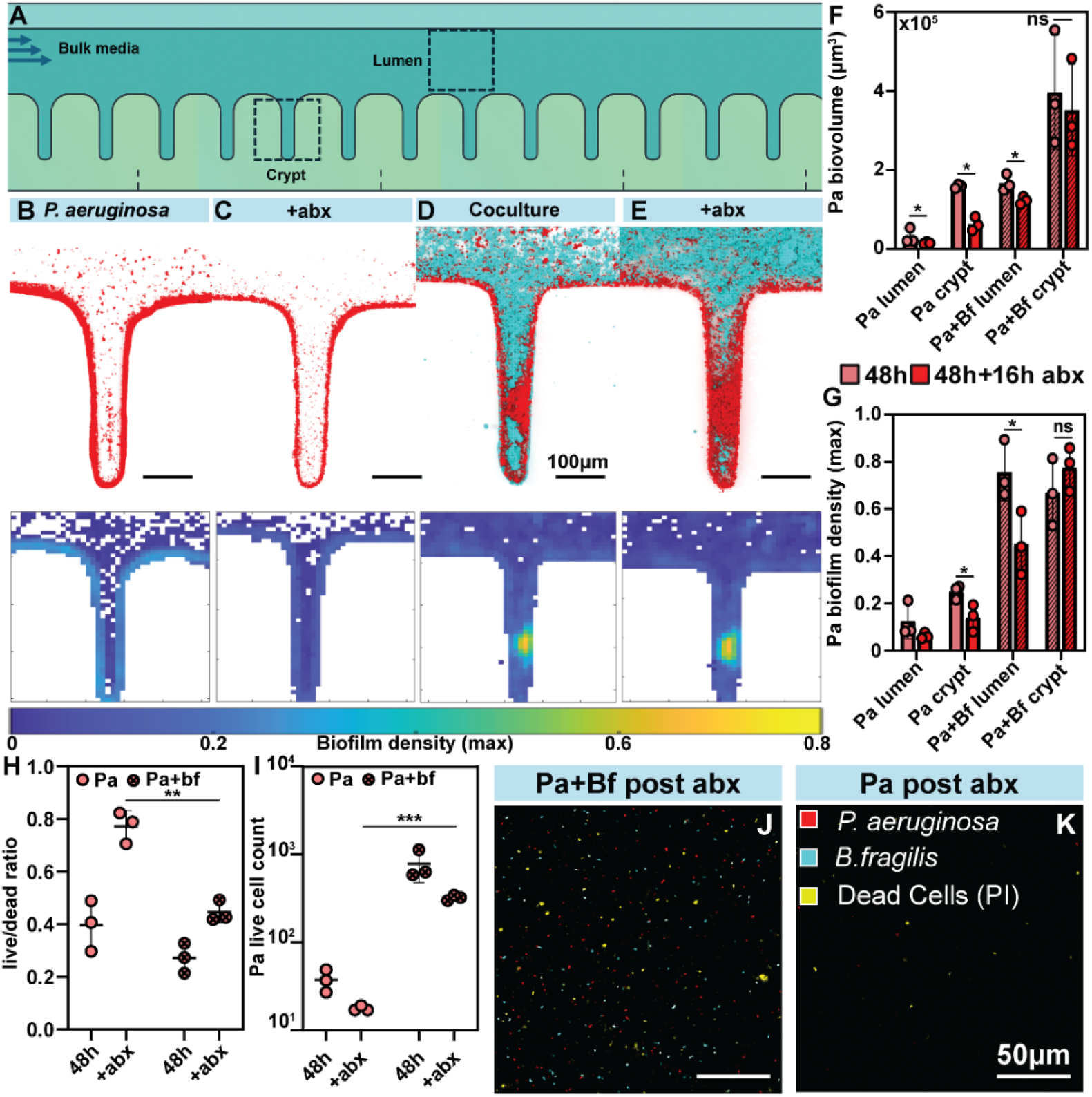
Gut architecture drives interaction between *B. fragilis* and *P. aeruginosa.* (A) Microfluidic crypt map. To understand how the microenvironment of the gut impacted the interaction of these two bacteria, a microfluidic device was fabricated that included crypt structures. **(B-E) 3D confocal reveals tight association in crypts**. Confocal images of Coculture crypts and monoculture Pa crypts. Pa was labeled with mGFP (red) and Bf was labeled with mRFP (cyan). **(F/G)Pa density is highest in coculture crypts.** Heatmaps of biofilm packing density outlining dense Pa growth and survival in coculture. **(H/I) Pa Antibiotic survival relies on biofilm formation**.Cell counts from biofilms that were dislodged, vortexed and stained with propidium iodide to identify dead cells. Ratio of live to dead cells in image and total live cell count determined by fluorescence. n=3 (H). **(J/K) Representative images of live dead staining (Yellow)**. Quantification of total biovolume of Pa in crypt and lumen environments n=3, Welch’s test *P<.05,**P<.01,***P<.001,****P<.0001(A). Quantification of the maximum density achieved by Pa in differing gut environments n=3, Welch’s test *P<.05,**P<.01,***P<.001,****P<.0001 (B).

As expected for the media and anaerobic conditions used, the *Pseudomonas* monoculture grew rather poorly, sparsely colonizing the crypt wall preferentially over the lumen **(Fig. 3B/F)**. The co-culture with *Bacteroides*, however, increased the *Pseudomonas* biomass substantially, particularly in the crypts **(Fig. 3D/F, microscopy image Pa+Bf**). Computational image analysis shows that the *Pseudomonas* populations were very dense and occupied an extensive section of the crypt (**Fig. 3F**).

We then treated the systems with 4 μg/ml ciprofloxacin, a concentration several fold higher than the typical MIC. In monoculture, *Pseudomonas* was significantly reduced in both the lumen and crypt spaces (**Fig. 3C/F).** In the co-culture, however, the *Pseudomonas* survived better (**Fig. 3E/F**), which is consistent with the mouse model. Interestingly, most surviving *Pseudomonas* resided in the crypts, and formed structures of much higher density than when alone. These bacterial structures remained largely unperturbed by the introduction of antibiotics, maintaining their high density. In contrast, the surface-attached *Pseudomonas* lost most of its structural density in all other contexts. High-density growth occurred mostly in close spatial proximity to *Bacteroides* (**Fig. 3G**)

We also performed an invasion experiment in which *Bacteroides* was first introduced, cultured for 24 hours, and then *Pseudomonas* was introduced. Not only did *Pseudomonas* invade the crypts, but it also grew and survived ciprofloxacin in the same manner as in the simultaneous coculture (**Fig. S3**).

To further test the interaction, we investigated the impact of *Bacteroides* on *Pseudomonas* using two classic biofilm assays: straight-chamber flow cells and static-well biofilms. In the standard straight-chamber assay, without the more intricate gut-mimicking architecture, *Pseudomonas* alone could only form a monolayer, and this thin biofilm was completely ablated by ciprofloxacin (**Fig. S3**). *Bacteroides* co-culture, however, induced production of a thick biofilm with an intricate spatial structure, in which the inner layers are predominantly composed of *Pseudomonas*, and the upper layers show a mixture of *Pseudomonas* and *Bacteroides*. Ciprofloxacin eliminated large clusters of *Bacteroides*, but the underlying *Pseudomonas* population survived (**Fig. S3**).

To confirm that the remaining biomass after antibiotic treatment was indeed viable, we subjected the microfluidic devices to extremely high flow to dislodge remaining biomass. These clumps of cells were then vortexed vigorously to mechanically disrupt them, and then were stained with the dead-cell-marker propidium iodide. This staining confirmed that *P. aeruginosa* within biomass colocalized with *B. fragilis* survives ciprofloxacin treatment significantly better than alone, exhibiting both a higher total surviving cell number and a lower live/dead ratio (**fig 3 H/I**). Representative images show that while dead cells are present in coculture, the population of both species appears largely viable (**Fig 3J/K**).

### *Bacteroides* acts as an anaerobic primary degrader, while Pseudomonadota converge as facultative secondary utilizers

To define nutrient use across this guild, we ran profiled carbon source utilization in minimal media with a panel of 190 sources from BIOLOG PM1 and PM2A. We carried out a 24h assay on*B. fragilis* and *P. aeruginosa*, as well as, *E. coli* and *K. pneumoniae*, profiling all four strains anaerobically and the three Pseudomonadota taxa in parallel aerobic controls (**Fig. 4; Fig. S4**). At the substrate level, *Bacteroides* was distinguished by strong anaerobic growth across complex carbohydrates and polymers, including dextrin and the cyclodextrins, whereas the three Pseudomonadota showed weaker and more restricted use of these types of carbon source (**Fig. 4A**). This separation was not simply a difference in overall growth magnitude. Instead, the *Bacteroides* species showed superior growth on compounds that most directly match a primary-degrader role, while the other taxa remained concentrated in a narrower downstream-accessible niche.

**Figure 4.**
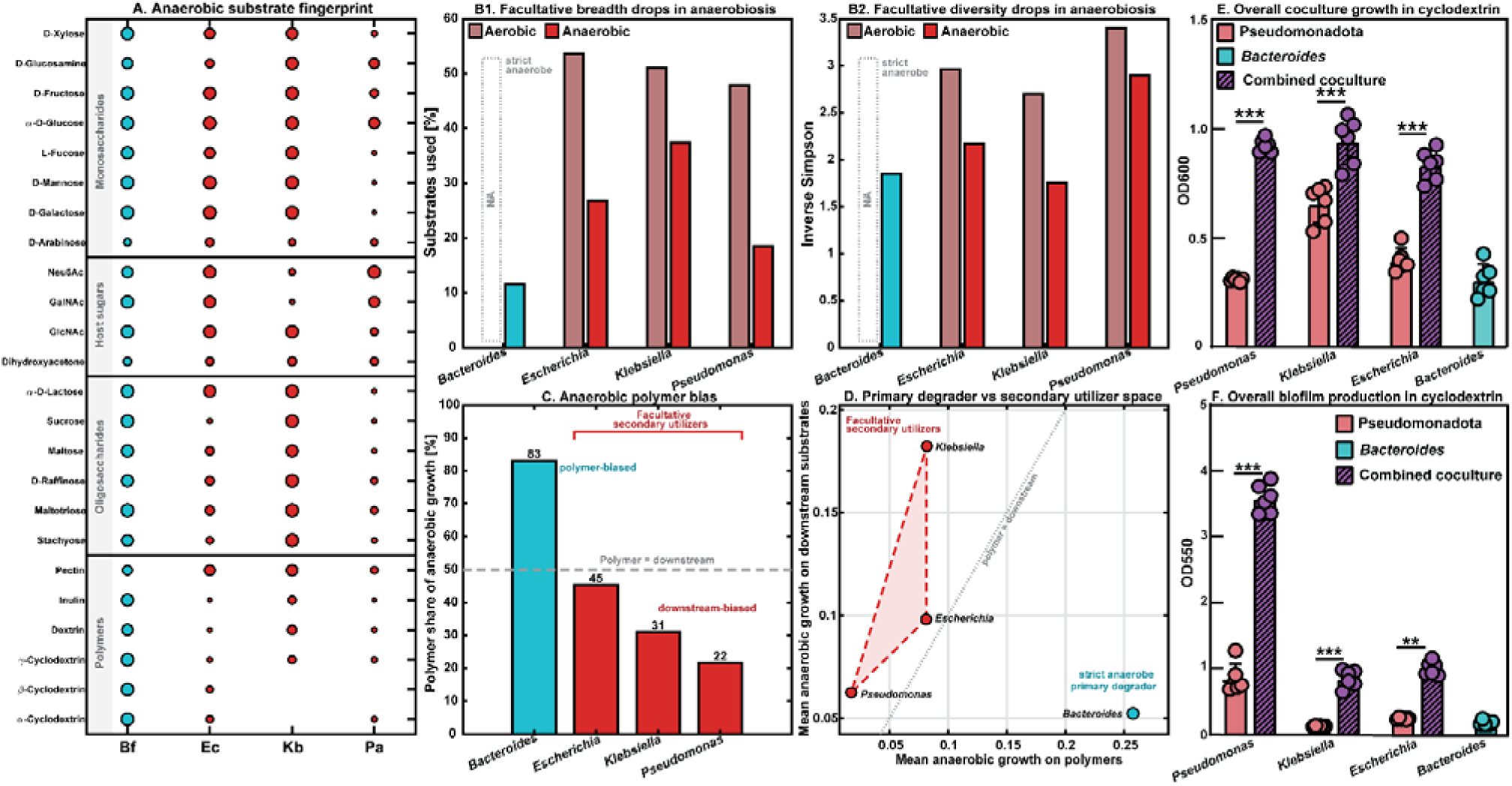
Substrate utilization patterns separate anaerobic *Bacteroides* from three Pseudomonadota and support a primary-degrader versus secondary-utilizer model. *Bacteroides* is anaerobic-only and polymer-biased, whereas *Escherichia*, *Klebsiella*, and *Pseudomonas* remain ecologically distinct from one another but converge as facultative secondary utilizers under anaerobic conditions. Cyan denotes *Bacteroides* and red denotes the Pseudomonadota throughout. **(A) Anaerobic substrate fingerprint.** Selected anaerobic responses from the 24 h BIOLOG PM1 and PM2A assays highlight the broad complex-carbohydrate and polymer profile of *Bacteroides* relative to the three Pseudomonadota; the full corrected nutrient map is shown in Supplementary Fig. 4C. **(B1) Facultative breadth drops in anaerobiosis.** Paired condition bars compare the breadth of substrate use across matched aerobic and anaerobic profiles for the three facultative Pseudomonadota. *Bacteroides* was assayed anaerobically only. **(B2) Substrate utilization diversity of Pseudomonadota drops in anaerobiosis.** Anaerobiosis reduces the diversity of nutrient classes used by the three facultative Pseudomonadota relative to aerobic controls. **(C) *Bacteroides* falls in the polymer-biased regime, whereas the Pseudomonadota occupy the downstream-biased regime.** Polymer bias compares mean anaerobic growth on polymers versus downstream substrates (Methods). **(D) *Bacteroides* is a primary degrader whereas Pseudomonadota are secondary utilizers.** The trophic-space plot uses the same polymer and downstream growth axes summarized in panel C. The shaded red polygon groups *Escherichia*, *Klebsiella*, and *Pseudomonas* as facultative secondary utilizers, while the cyan *Bacteroides* point marks a strict-anaerobe primary degrader position. **(E/F) Growth and biofilm output increase in direct coculture**. Coculture assays of *B. fragilis* with Pseudomonadota show growth increase N=6, Welch’s test ***P<.001 (E). Crystal violet staining was then performed on the same wells N=6, Welch’s test **P<.01,***P<.001(F).

Anaerobiosis reduced the breadth and diversity of substrate use across the three facultative Pseudomonadota: Relative to aerobic growth, each taxon showed a contraction in both the fraction of substrates supporting growth (**Fig. 4B1**) and the diversity of nutrient classes among those substrates (**Fig. 4B2**). *Bacteroides* was represented only in anaerobiosis in these comparisons because it is a strict anaerobe, emphasizing that the contrast is between an anaerobe specialized for this environment and facultative anaerobes whose metabolic breadth is limited within it.

Despite taxon-specific differences in the downstream substrates they favored, *Escherichia, Klebsiella*, and *Pseudomonas* still formed a shared ecological grouping relative to *Bacteroides*. In a polymer-bias score, *Bacteroides* alone occupied the polymer-biased regime, whereas the three Pseudomonadota remained downstream-biased (**Fig. 4C**). The same separation was evident in trophic space, where *Escherichia, Klebsiella*, and *Pseudomonas* located in the region characteristic of secondary utilizers and *Bacteroides* located in a distinct primary-degrader region (**Fig. 4D**). Thus, the key commonality among the Pseudomonadota was not identical substrate preference, but a shared ecological deficit in their ability to use polymers anaerobically.

Together, these data support a model in which *Bacteroides* is a strict-anaerobe primary degrader, while diverse Pseudomonadota are facultative secondary utilizers, limited to sharing a downstream niche.

Consistent with this model, when co-cultured with *B. fragilis* on cyclodextrin as the sole carbon source under anaerobic conditions, all three Pseudomonadota species exhibited enhanced growth (**Fig. 4E**).

This metabiotic support had broader phenotypic consequences: co-culture with *Bacteroides* increased anaerobic biofilm formation in cyclodextrin medium and rendered these mixed-species biofilms more resilient to ciprofloxacin treatment (**Fig.4F;Fig S5**). Together, these results indicate that *Bacteroides* polysaccharide breakdown can expand the accessible nutrient landscape for diverse Pseudomonadota in anaerobic environments, a metabiotic process that promotes both colonization and antibiotic tolerance.

### Metabiosis potential varies among *Bacteroides* species

To test whether Pseudomonadota can exploit metabiotic degradation of complex polysaccharides by *Bacteroides*, we quantified growth in liquid co-cultures of *P. aeruginosa* and *Bacteroides fragilis* with cyclodextrin as the sole carbon source. The co-culture entered exponential growth substantially earlier than either monoculture (**Fig. 5A**), consistent with the idea that *Bacteroides* generates diffusible resources that accelerate Pseudomonadota growth.

**Figure 5:**
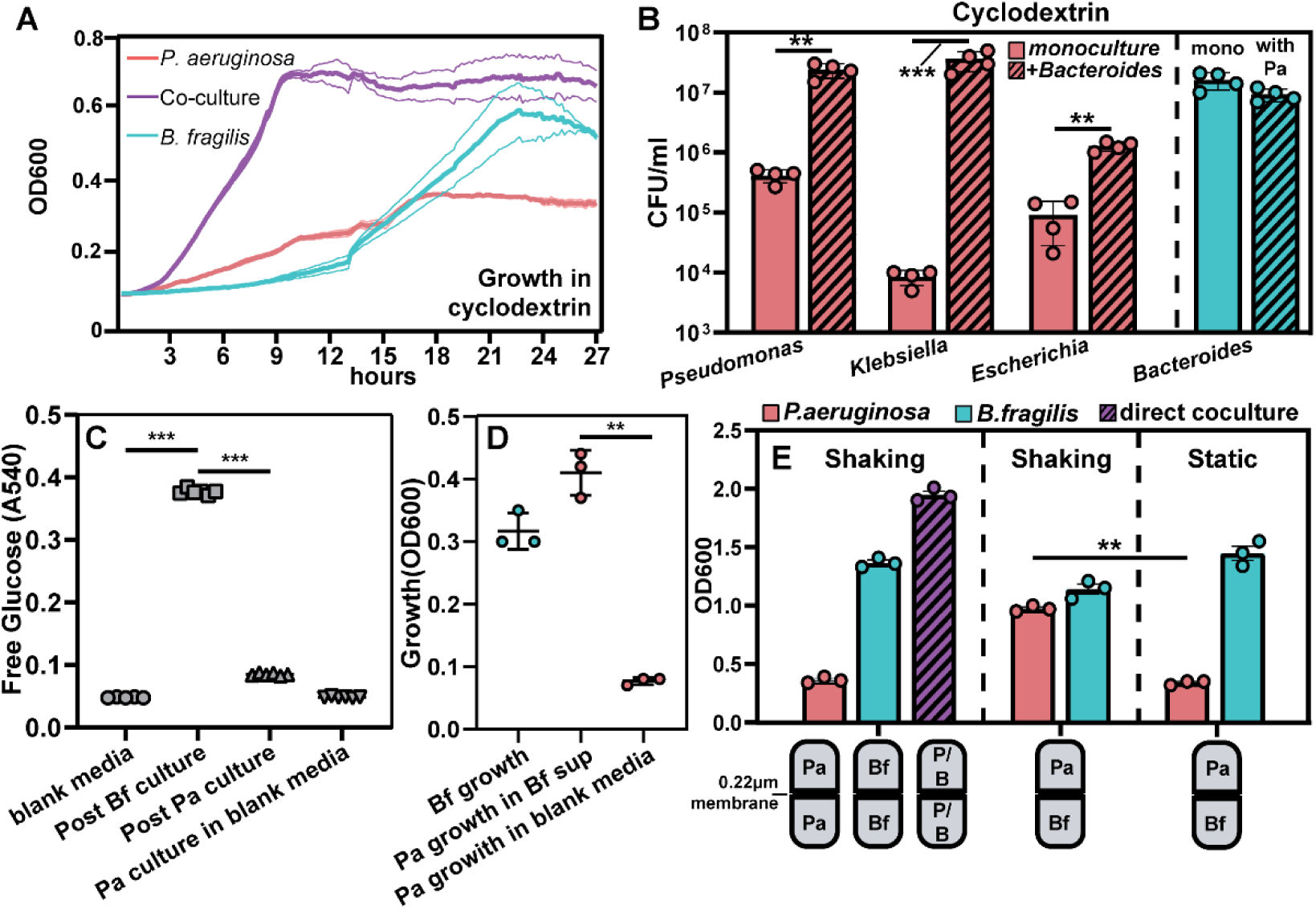
*B. fragilis* breakdown of soluble fiber is a public good that Pseudomonadota can utilize. **(A) coculture in cyclodextrin yields early growth.** A growth curve of Pa and Bf in cyclodextrin outlining the dynamics of coculture growth. **(B) Cyclodextrin coculture explicitly benefits Pseudomonadota**. CFU counts for each Pseudomonadota pathogen outlining the growth benefit of coculture with Bf in cyclodextrin N=4, Welch’s test **P<.01,***P<.001 (B**).(C/D) Glucose is released by Bacteroides and Consumed by Peudomonadota**. Supernatant experiments assaying for free glucose in spent media and the final growth amounts that match N=6 (C) N=3 (D), Welch’s test **P<.01,***P<.001 (C-D). **(E) Local association is required for growth benefit**. Cerillo duet culture assays showing that the growth benefit with Bf is diffusible and distance dependent N=3, Welch’s test **P<.01 (E).

This metabiosis generalized beyond *P. aeruginosa*: both *K. pneumoniae* and *E. coli* exhibited analogous co-culture growth enhancement with *B. fragilis* (**Fig. S5**). Quantitative plating further showed that *Bacteroides* can amplify the Pseudomonadota population (by up to ∼10,000-fold for *K. pneumoniae*) without detectably altering *Bacteroides* abundance (**Fig. 5B**).

To determine whether this metabiotic interaction reflects the public release of monosaccharides during polysaccharide degradation, we grew *P. aeruginosa* in cell-free spent media collected from *B. fragilis* cultures grown on cyclodextrin (a glucose polymer) or inulin (a fructan). Although *B. fragilis* can easily utilize both substrates, Pseudomonadota cannot (**Fig. S5**). Nonetheless, all species of Pseudomonadota grew robustly in spent media from either condition (**Fig. S5**). Consistent with monosaccharide release, spent media contained free glucose or fructose, respectively, and these sugars were depleted during *P. aeruginosa* growth (**Fig. 5C–D; Fig. S5**).

Next, motivated by the close proximity between *Pseudomonas* and *Bacteroides* observed in the crypt model (**Fig. 3D; Fig. S5**), we asked whether spatial separation limits access to these breakdown products. Using a two-chamber co-culture system separated by a 0.22μm membrane that prevents cell contact while allowing diffusion, we found that *B. fragilis* supported *P. aeruginosa* growth both when co-resident in the same chamber and when confined to the opposite chamber under shaking conditions (**Fig. 5E**). Under static conditions, where transport across the membrane is expected to be slower, *P. aeruginosa* growth was reduced (**Fig. 5E**). This supports the notion that passive diffusion of *Bacteroides*-derived breakdown products underlies the interaction and suggests that close proximity can enhance exploitation in structured environments.

Finally, because *Bacteroides* species vary in their polysaccharide-degrading repertoires, we tested whether the metabiosis depended on their capacity to utilize specific polysaccharides. Prior work has shown that different *Bacteroides* encode distinct polysaccharide utilization loci (PULs) and therefore differ in their ability to degrade substrates such as inulin^35^. Consistent with this prediction, co-culture of *P. aeruginosa* with *Bacteroides ovatus* (an inulin utilizer) supported metabiotic growth on inulin (**Fig. S5**), whereas co-culture with *Phocaeicola vulgatus,* formerly *Bacteroides vulgatus*, (which lacks key inulin-utilization determinants) did not (**Fig. S5**). Together, these results show that Pseudomonadota can benefit from the metabiotic polysaccharide degradation through diffusible monosaccharides, and that this interaction is regulated by spatial transport and by the commensal-specific polysaccharide utilization capacity.

### Metabiotic polysaccharide breakdown underlying microbiota permissiveness in allo-HCT

Guided by the species-conditional mechanism, we returned to the clinical cohort to ask whether metabiotic potential through polysaccharide breakdown is encoded in the post-prophylaxis microbiota and could help explain, and anticipate, permissiveness to Pseudomonadota expansion. We assembled patient fecal metagenomes into high-quality MAGs and quantified “metabiotic polysaccharide degradation potential” as the number of digestive carbohydarate-active enzyme (CAZyme) gene clusters (CGCs) that include at least one secreted digestive CAZyme (signal peptide–positive), reasoning that extracellular/periplasmic enzymatic steps generate diffusible breakdown products that can be exploited by neighboring cells.

At the species level, the MAGs of *Bacteroides* and its close relatives, *Phocaeicola,* showed substantial heterogeneity in the predicted substrates and total counts of these secreted digestive CGCs (**Fig. 6A**), indicating that the capacity to generate publicly accessible glycan breakdown products varies across the genomes of primary degraders in patient microbiomes. We then broadened this analysis across the major lineages identified in the patient cohort. The Bacteroidota phylum, and in particular the families Bacteroidaceae and Tannerellaceae, stood out as uniquely enriched for CGCs with signal peptides (**Fig. 6B**), consistent with these commensals being dominant contributors to the metabiotic polysaccharide degradation potential in the gut ecosystem. Importantly, despite Tannerellaceae sharing high predicted degradation potential, Bacteroidaceae were significantly more abundant and ∼four times more prevalent than Tannerellaceae in the post-quinolone-prophylaxis window (**Fig. 6C**), making Bacteroidaceae the most plausible and widespread “producer” lineage capable of shaping glycan accessibility after prophylaxis start.

**Figure 6.**
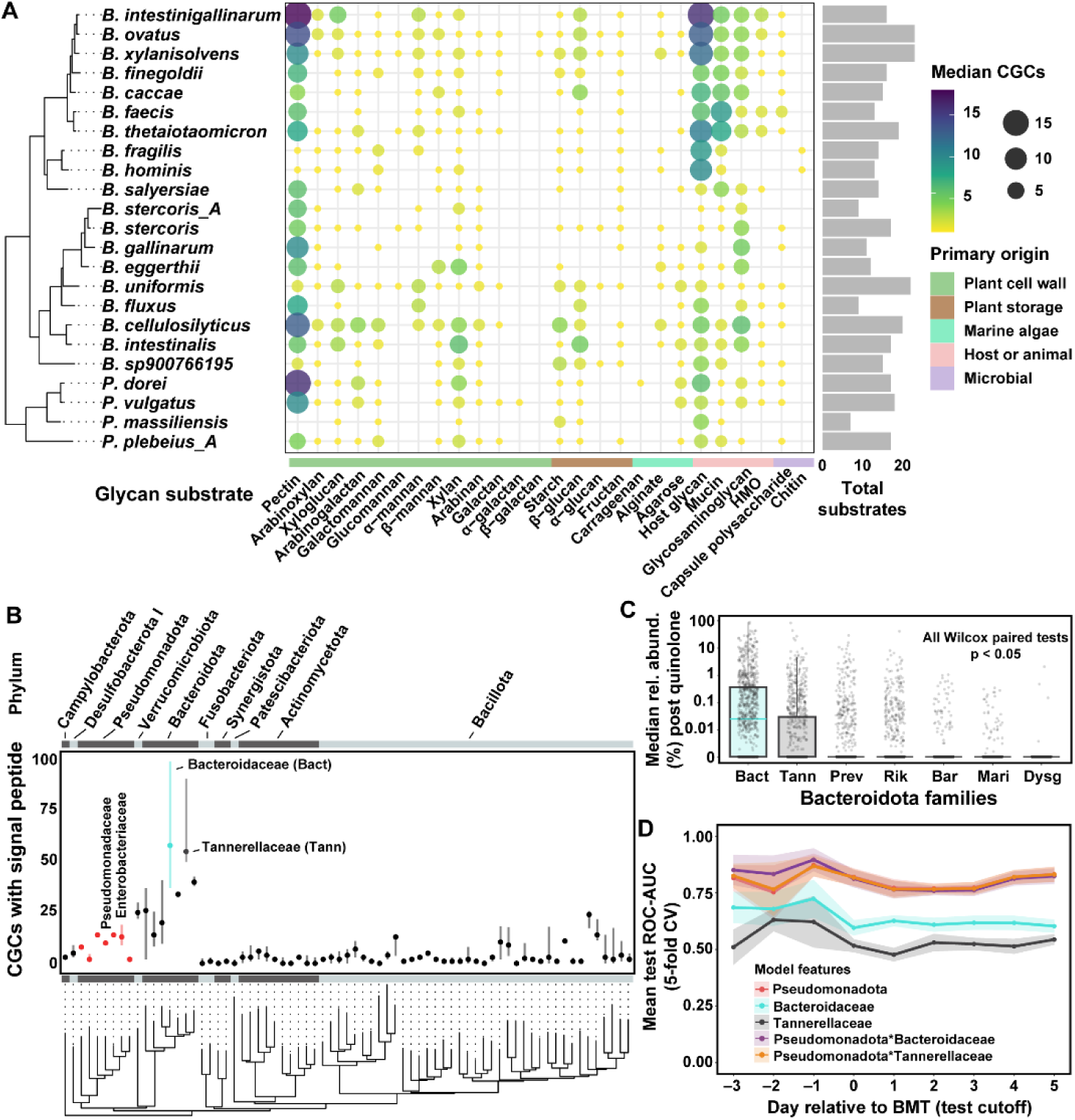
Metagenomics links commensal glycan-processing potential to microbiota permissiveness and early prediction of Pseudomonadota-associated bacteremia in allo-HCT. (A) Genomic potential for Metabiotic glycan degradation varies across common Bacteroidaceae species reconstructed as metagenome-assembled genomes (MAGs). Bubble size and color denote the median number of predicted digestive CAZyme gene clusters (CGCs) targeting each glycan substrate, with substrates grouped by primary origin (plant cell wall, plant storage, marine algae, host/animal, microbial). Bar plot summarizes the total number of glycan substrates with detectable CGCs per species. **(B) Cohort-wide comparison of predicted metabiotic glycan degradation potential across major gut lineages.** For each lineage (phylogenetically ordered), points indicate the total number of CGCs containing a signal peptide (secreted digestive capacity), highlighting the uniquely high values in Bacteroidota—particularly Bacteroidaceae and Tannerellaceae—relative to other dominant phyla (including Bacillota) and pathogen-associated lineages (Pseudomonadota/Enterobacteriaceae). **(C) Post-quinolone prophylaxis**, Bacteroidaceae are significantly more abundant and more prevalent than Tannerellaceae and other Bacteroidota families, positioning them as the dominant putative glycan “producers” during the permissive window (paired Wilcoxon tests, all *p* < 0.05). **(D) Early risk modeling:** mean test ROC-AUC (5-fold cross-validation) for logistic regression models predicting subsequent Gram-negative bacteremia as a function of day relative to BMT, using Pseudomonadota abundance alone or combined with commensal family features. Incorporating Bacteroidaceae improves discrimination relative to Tannerellaceae and provides incremental signal beyond pathogen abundance alone early in the peri-transplant course.

Finally, we asked whether these commensal features provide an actionable early signal by fitting simple logistic regression models to predict Gram-negative bloodstream infection during neutropenia, using per-patient median relative abundances accumulated longitudinally over time (stratified 5-fold CV).

Consistent with the cohort ecology, models using Bacteroidaceae features yielded a modest improvement in early discrimination relative to Tannerellaceae features (**Fig. 6D**), supporting the idea that metagenomic/phylogenetic readouts of PUL/CGC-encoded producer capacity, combined with the post-antibiotic community context, can help quantify microbiota permissiveness before overt pathogen expansion.

## Discussion

In healthy individuals, the human gut microbiota is dominated by commensal bacteria from the phyla Bacillota (formerly Firmicutes) and Bacteroidota (formerly Bacteroidetes). The ratio between these two phyla (the Firmicutes/Bacteroidetes ratio) has been widely discussed as a marker of gut-associated disease states^36^, perhaps because it is an approximation for the ecological balance between primary utilizers and downstream consumers, and it is easy to imagine how dysregulation could arise when these linked trophic roles become uncoupled. However, a major source of clinical harm, especially in immunocompromised patients, comes from a minority phylum, Pseudomonadota. This phylum is typically low in abundance but contains many pathogenic species that drive a large fraction of severe infections^15^. In allo-HCT, in particular, these bacteria can exploit ecological disturbance into invasive disease. Our results show that their emergence is not simply a dysbiosis, but a predictable consequence of how antibiotic perturbation exposes resource producers within the remnant commensal population.

For our work, we leveraged a large-scale allo-HCT cohort to provide an extreme yet quantifiable setting in which the microbiota undergoes ecological disturbances with consequences for clinical outcome^29,37^. Previous work has shown that fluoroquinolone prophylaxis reduces but does not eliminate these bloodstream infections, and that risk concentrates in narrow windows, during which domination and bacteremia surge during patient neutropenia^15^. However, Pseudomonadota dynamics are fast and can be easily missed without daily sampling. This limits the use of surveilling Pseudomonadota relative abundances as an actionable early warning system. We demonstrate that polysaccharide degradation by gut commensals creates a metabiotic niche that is normally suppressed by competitive exclusion from resident secondary consumers, but becomes accessible to Pseudomonadota following antibiotic-driven competitive release^38,39^. Metabiosis, in which one organism’s metabolic activity passively generates conditions that benefit a subsequent colonizer, has been described in successional ecology and human health^23,40^. Here we show it operates as a clinically consequential dynamic in the antibiotic-damaged gut. The persistence of *Bacteroides* driven carbohydrate processing capacity after prophylaxis thus creates an open niche that supports Pseudomonadota growth, biofilm formation, and antibiotic tolerance under anaerobic conditions.

Members of the Pseudomonadota vary in their metabolic abilities, but they share a common ecological constraint in the anaerobic, polymer-rich gut. Since most are inefficient at degrading complex polysaccharides, they rely on metabolites produced by primary degraders such as *Bacteroides*. They all act as secondary consumers of public goods, despite differences in downstream metabolism. In our work, we used *P. aeruginosa* as the main model to study this dependency, not as a universal proxy for all gut Pseudomonadota. The signal in patient data is clearest at the phylum level; genus-and species-level differences affect infection dynamics but reduce predictive power when resolved to finer levels. By focusing on this ecologically constrained group of facultative anaerobic Gram-negative pathogens, metabiotic polysaccharide breakdown reveals a principle applicable across many infections, unlike interactions limited to specific taxa or nutrients. Previous work has shown that antibiotics can increase access to mucosal carbohydrates, which, in turn, facilitates pathogen expansion^19^. Ecosystem collapse and recovery after antibiotic use can also be shaped by diet, with a low-fiber diet providing less substrate for polysaccharide breakdown, aggravating collapse and delaying recovery^41^. Our data show that in allo-HCT, the persistence of microbiota-driven carbohydrate-processing capacity after prophylaxis can enable Pseudomonadota expansion and predict downstream bloodstream infection in a clinical setting.

This view extends classic post-antibiotic expansion paradigms beyond host-derived carbohydrates to community-derived public goods with direct clinical consequences and suggests that shared biological constraints can provide broader targets for intervention.

A broader translational implication is that mechanistic ecology can be used to improve prediction and pinpoint intervention targets. Instead of waiting for pathogen detection or dominance, we suggest based on our mechanistic insights and ecological profiling that high-risk states can be identified earlier, allowing for preventative measures while still feasible. This perspective also clarifies why targeted interventions may not generalize across pathogen classes: for example, a commensal consortium strategy that decolonizes Enterobacteriaceae by controlling gluconate availability is specific to the impact of a particular nutrient on a specific taxon^42^. Our work highlights an alternative axis in which the metabiotic potential of polysaccharide breakdown remaining in antibiotic-damaged microbiota can support secondary degraders as a broad and tractable niche feature.

Finally, this study exemplifies the concept of reverse translational research^43^. We started with human, time-resolved prospective observations; use reductionist mouse and *in vitro* gut-mimicking systems to establish a causal mechanism; and then return to the clinical population to develop mechanism-informed insights from patient microbiome profiling. In a disease context where hours to days can separate containment from lethal bloodstream infection, reverse-translational insights can accelerate our ability to turn microbiome ecology into actionable medicine.

## Methods

### Media, bacterial strains and culture conditions

All bacterial strains were grown overnight prior to experiments in BHI supplemented with hemin. Pseudomonadota were incubated aerobically with shaking at 37 C and all *Bacteroides* and *Phocaeicola* species were grown at 37 C shaking inside a COY labs anaerobic chamber. *P. aeruginosa* strain PA01 and *B. fragilis* strain NCTC9343 had chromosomally integrated fluorescent protein expression of mGFP and mRFP, respectively (Table S1). For coculture, carbon source utilization, and BIOLOG assays, a *Bacteroides*-specific minimal medium supplemented with nitrate was used, as described previously^34^.

Importantly, the carbon source of interest, e.g., cyclodextrin, inulin, was added at a concentration of 5mg/ml. Prior to usage, this media is placed in an anaerobic environment overnight to allow for degassing.

### Microfluidic community assays and device assembly

Both straight chamber and gut mimic microfluidic devices were cast from PDMS using standard soft lithography techniques^44^. Briefly, 20ml of polymer is poured into the cast and placed under vacuum to eliminate bubbles. Once the polymer is clear, it is left to cure overnight at 37C. The following day, a scalpel is used to remove the polymer cast and divide the devices. Each device is then exposed to air plasma and bonded to a #1.5 glass coverslip. Once assembled, inlet and outlet tubing is placed into the device. This PTFE tubing has an internal diameter of 0.3mm and can be placed over a 25 gauge needle. This needle is then affixed to a BD 1ml plastic syringe filled with *Bacteroides* minimal media. These syringes are then placed on a Harvard apparatus Pico plus syringe pump.

To inoculate the devices for bacterial coculture, each overnight culture was normalized to an OD600 of 1.0. All of the equipment and devices are then passed into the anaerobic chamber. Each species was then mixed in equal part and 100μl of culture was passed into the device using a standard micropipette. The tubing is then placed into the device and placed into the incubator set to 37C. The devices are left to incubate for 30 min to allow for initial attachment. After this phase, the pump is turned on at a flow rate of 0.1μl/min. For the introduction of antibiotics, a 4 µg/ml ciprofloxacin solution was prepared in *Bacteroides* minimal medium. After 48 hours of initial biofilm growth, the media syringe is swapped and media containing antibiotic is allowed to perfuse overnight after which imaging was performed. In the invasion experiment, *Bacteroides* was first inoculated into the device and allowed to grow for 24 hours. After this, a media syringe containing *Bacteroides* minimal media and OD600 0.1 *P. aeruginosa* was swapped and allowed to flow for 1 hour. Afterwards, the media syringe was swapped again to sterile *Bacteroides* minimal media. Antibiotic exposure for these experiments was carried out in the same fashion as above.

To dislodge biomass after the experiment, the flow rate was placed on its maximum setting on the pump and allowed to effuse into a 1.5 ml micro centrifuge tube. This effluent was then vortexed for several minutes, and pipette mixed repeatedly to eliminate cell clusters. 100μl of this effluent was then stained with 1μl of 20mM propidium iodide and allowed to sit for 2 minutes. 10μl from each condition was then spotted onto a glass slide with a cover slip and imaged.

### Coculture conditions, biofilm assays

Coculture assays were carried out in 96 well format. After overnight culture, each species was normalized to OD600 1.0 in *Bacteroides* minimal medium. Each normalized culture was then inoculated into fresh minimal medium to reach a final OD600 of 0.05. 200μl of culture was then placed into individual wells and left to incubate within the anaerobic chamber statically overnight. For growth curves, 96 well plate cultures were placed into a Tecan Sunrise plate reader within the anaerobic chamber. Plates were shaken and read at 15 minute intervals. After overnight growth, plates were removed from the anaerobic chamber and OD600 was measured. If CFU counts were desired, a 20μl aliquot was taken from each culture and serially diluted in PBS. 5ul from each serial dilution was then spotted onto sheep’s blood agar in triplicate and then left to incubate anaerobically overnight. Colonies of each species were then counted and differentiated by colony morphology

To measure the biofilm formation of each well culture, a standard crystal violet assay was used. All supernatant was removed from the well and 200μl of 0.1% crystal violet was pipetted in. This solution was left to stain for 5 minutes. Afterward, the crystal violet is removed and the stained biofilms are washed with 200μl PBS. After washing the stained biomass is dissolved using 30% glacial acetic acid and pipetted several times to mix. These wells can then be quantified using a Tecan Spark plate reader at an absorbance of OD550 Minimum inhibitory concentrations were determined using standard methods. Bacterial culture was inoculated into *Bacteroides* minimal medium at a final OD of 0.05. Serial dilutions of ciprofloxacin were then placed into each well decreasing by 2 fold. These plates were then incubated anaerobically overnight and MIC was determined visually.

### BIOLOG assays

PM1 and PM2a plates were purchased from BIOLOG. These plates contain 1 individual carbon source per well. To prepare cells for BIOLOG, overnight cultures are taken and normalized to an OD600 of 1.0. 1ml of culture is then placed within a 1.5ml micro centrifuge tube and spun at 10,000 rpm within a microcentrifuge for 2 minutes. After this spin, the supernatant is removed and cells are resuspended in 1ml fresh, carbon free *Bacteroides* minimal medium. This spin was then repeated 2 additional times to eliminate any residual carbon from the media. After washing, these cells are then inoculated into fresh *Bacteroides* minimal media to reach a final OD600 of 0.05. 150ul of inoculum is then placed into each well of the BIOLOG plate. This media must be pipetted up and down several times to make sure that the carbon sources are suspended and equally mixed within the well. Once all carbon sources are suspended, each BIOLOG plate was placed in the anaerobic chamber incubator at 37C and allowed to grow overnight. After incubation, plates were removed from the anaerobic chamber and OD600 was measured using a Tecan spark plate reader.

For each strain-condition profile, PM1 wells (1-96) and PM2A wells (97-192) were background-corrected separately. Background for each plate was estimated as the median of the 10 lowest well OD values, and that plate-specific background was subtracted from all wells on the same plate. Broad nutrient annotations were collapsed to Control, Polymer, Carbohydrate, Amino acid, Carboxylic acid, and Other. Raw and corrected well-level assay distributions are shown in Supplementary Fig. 4A-B.

For nutrient maps and mean-growth summaries, negative background-corrected values were truncated to 0. Breadth in Fig. 4B1 was defined as the percentage of non-control wells with background-corrected OD > 0.1. Diversity in Fig. 4B2 was computed as an inverse Simpson index across the broad nutrient classes represented among substrates exceeding the same 0.1 utilization threshold. Polymer bias in Fig. 4C was calculated as 100 x polymer / (polymer + downstream), where polymer is the mean anaerobic background-corrected OD across polymer wells and downstream is the mean anaerobic background-corrected OD across non-polymer, non-control wells after truncation of negative values to 0. Fig. 4D uses these same two mean-growth quantities as the x-axis (polymer) and y-axis (downstream) coordinates. Fig. 4A shows a selected anaerobic substrate fingerprint drawn from the full corrected nutrient map in Supplementary Fig. 4C. In Supplementary Fig. 4C, nutrients were ordered first by broad class and then by mean corrected OD across all strain-condition profiles. Bubble size in Fig. 4A and Supplementary Fig. 4C scales with asinh(corrected OD / 0.05). Colors follow the manuscript palette: cyan for anaerobic *Bacteroides*, bright red for anaerobic Pseudomonadota, and dim red for aerobic Pseudomonadota.

### P. aeruginosa supernatant toxicity

*P. aeruginosa* was cultured in *Bacteroides* minimal media supplemented with 5mg/ml glucose in either aerobic or anaerobic conditions overnight at 37C. 1ml of overnight culture was then spun down in a centrifuge at 10,000rpm and supernatant was removed. This supernatant was then filter sterilized through 2 sets of 0.22μm syringe filters to achieve cell free supernatant. This supernatant was then used to culture *Bacteroides* at a starting OD of 0.05. Supernatant was serially diluted into fresh *Bacteroides* minimal media with 5mg/ml glucose. For the no supernatant control. *Bacteroides* minimal media with no carbon source was serially diluted with fresh media containing 5 mg/ml glucose. All conditions were the cultured anaerobically overnight.

### Supernatant monosaccharide assay

To determine breakdown of polysaccharide and release of free glucose, several cultures and supernatants were used. First, *Bacteroides* and gamma Pseudomonadota were prepared in the same fashion as a standard coculture assay. These initial bacterial cultures were then inoculated alone in complex polymer and allowed to grow anaerobically overnight. After incubation, cultures were measured for growth (OD600). These cultures were then spun down at 10,000 rpm to separate cells. The supernatant was then removed and passed 2 times through a 0.22μm sterile syringe filter. A small aliquot of these supernatants was taken and mixed with a commercially available glucose/fructose assay kit. Free glucose and fructose were determined by absorbance at 650 and 340 respectively. The remainder of the *Bacteroides* supernatant was then reinoculated with *P. aeruginosa* to a final OD of 0.05 and allowed to incubate anaerobically overnight. After this second incubation, growth was measured and cells were removed to measure free monosaccharide as stated previously.

Alto plate reader and Duet coculture system were purchased from Cerillo. These duet cuvettes have two chambers separated by a 0.22μm filter. For these assays, bacterial inoculum was prepared as stated for coculture assays above, except each species was inoculated on opposite sides of the 0.22μm filter. Each side of the cuvette was filled with 500ul of BMM and inoculated to a final OD600 of 0.05. These cuvettes were then placed into an Alto plate reader which was then deposited into the anaerobic chamber and left to incubate in either shaking or static culture. The Alto plate reader then collected OD600 data, which could then be read and interpreted by Cerillo Labrador software.

### Mouse model and FISH staining

Female C57BL/6J mice, aged 6–8 weeks, were single housed in autoclaved cages throughout the experiment. Animals received autoclaved drinking water supplemented with a cocktail of streptomycin (2 mg/ml) and penicillin (1,500 U/ml) for one week. The antibiotic solution was replaced once, three days after the initiation of treatment. Irradiated 5053 chow was provided ad libitum during this period.

Following cessation of antibiotics, each mouse was orally gavaged with approximately 107 CFUs of either Pseudomonas aeruginosa, *Bacteroides fragilis*, or a 1:1 mix of both strains in PBS. Fecal samples were collected daily during first week post-gavage, then once weekly during the second and third weeks from each mouse. Samples were plated on Cetrimide agar (selective for *P. aeruginosa*) and Sheep Blood Agar (selective for *B. fragilis*) to monitor colonization levels. Once stable colonization was established (14–21 days post-gavage), each animal was weighed and administered ciprofloxacin intraperitoneally at a dose of 25 mg/kg body weight for five consecutive days. Mice were monitored, fecal samples were collected daily during first week post-ciprofloxacin, then one-two times weekly for up to 3-4 weeks and plated on selective media to assess the impact of ciprofloxacin on bacterial colonization.

At the end of the experiment, colon sections were harvested from the mice gavaged with both Pa and Bf. These sections were then fixed in formalin and then sliced via microtomy. These slices were then affixed to slides and stained with 2 FISH probes, one specific to *P. aeruginosa* 489158 (B-*Pseudomonas*-23S-degen), and the other *B. fragilis* 843618-C2 *(B. fragilis)*.

### Clinical Data Sources and Input Tables

Analyses were recomputed from raw clinical metadata, microbiome count tables, taxonomy annotations, and infection records published in^16^. Inputs comprised longitudinal sample metadata, infection metadata, peri-transplant drug exposures, taxonomy assignments, and ASV-level count tables covering the allo-HCT peri-transplant interval.

### Episode Reconstruction and HCT Anchoring

Episodes were reconstructed from sample-linked HCT anchors using EpisodeID = PatientID HCTTP_(Timepoint DayRelativeToNearestHCT), with values rounded to six decimal places to stabilize joins across tables. Sample metadata were restricted to rows with non-missing SampleID, non-missing PatientID, finite Timepoint, and finite DayRelativeToNearestHCT, and day values were rounded to integer day-relative coordinates (DayRelInt). The resulting episode table was linked back to HCT metadata by the same HCT anchor.

This reconstruction produced the full descriptive cohort of 1318 transplant episodes in 1276 patients.

### First Gram-negative Bloodstream Infection Definition and Exclusions

First bloodstream infection events were taken from the infection metadata after restricting to rows with non-missing PatientID, non-missing infectious agent, finite event timing, and day-relative position in the interval day-15 to +30.

Infections labeled as Enterococcus or Streptococcus were excluded before defining the Gram-negative bloodstream infection endpoint. Within each episode, the first remaining qualifying event in the day-15 to +30 window was retained as the first Gram-negative bloodstream infection. This yielded 99 first Gram-negative bloodstream infection episodes in the full cohort. For modeled BSI start-stop analyses, only first events with EventDay >=-15 and EventDay <= 30 were coded as inferential events, which yielded 64 modeled BSI events in the leakage-safe inferential denominator.

### Prophylaxis Start Definition

Prophylaxis start was defined episode-wise from the drug table as the earliest quinolone row active on day-2 relative to transplant. Drug rows were first episode-linked using their start and stop timepoints relative to transplant, then restricted to quinolone-category rows spanning day-2. For each episode, PrimaryPostStartDay was set to the minimum start day among qualifying quinolone rows. If no such qualifying quinolone row was available, prophylaxis start was set by default to day-2. This definition was used throughout all post-prophylaxis and inferential analyses.

### Taxonomic Aggregation and Residual-community Construction

ASV counts were joined to taxonomy after normalizing phylum, family, and genus labels. Pseudomonadota and legacy Proteobacteria labels were treated as analytically equivalent and grouped together when defining the Pseudomonadota burden. Residual-community analyses removed all Pseudomonadota/Proteobacteria reads from each sample and renormalized the remainder to unit relative abundance. Residual Bacteroides was then defined as the renormalized relative abundance of Bacteroides within that residual community. log-scale transforms used a fixed pseudocount of 1e-5 or 1e-6. Residual-community features also included inverse residual Simpson diversity, residual Bacteroidota-to-Bacillota balance, and prevalence-ranked residual families.

### Full Cohort and Inferential Denominator Construction

The full descriptive cohort included all reconstructed episodes with valid episode identity, regardless of post-prophylaxis follow-up sufficiency. The inferential denominator was then restricted to episodes with at least one eligible post-prophylaxis observation in the modeled day-15 to +30 window before any qualifying Gram-negative bloodstream infection event. This leakage-safe subset contained 1052 episodes in 1027 patients. For descriptive panels, timing was displayed on the transplant-relative clinical clock. For inferential panels, analyses used one-day start-stop intervals on a separate inferential clock spanning the same biological day range.

### Metagenome assembly, and binning

We trimmed raw reads with --cut_mean_quality = 20 and --length_required = 30 using fastp v0.24.1.^45^ Then, we assembled the trimmed reads into contigs using metaSPAdes v4.1.0^46,47^ with k-mer lengths 21, 33, 55, 75, and 95. To bin the assembled contigs into metagenome-assembled genomes (MAGs), we employed a multi-sample binning strategy wherein sequencing depth is estimated by mapping reads from every metagenomic sample of a given patient to the contigs assembled from all the samples of the same patient.^48^ This increases binning efficiency and accuracy^48,49^. We mapped reads using strobealign v0.16.0^50^ and binned contigs using VAMB v5.0.4^51^, considering contigs ≥ 1000 bp and keeping bins ≥ 250,000 bp. Next, we used CheckM2 v1.1.0^52^ to obtain MAG-level assembly quality statistics and GTDB-Tk v2.4.1^53^ to assign MAG taxonomy. For subsequent analyses, we used MAGs with ≥ 90% completeness, ≤ 10% contamination, and contig N50 ≥ 50,000 bp. This resulted in 10,360 MAGs representing 980 bacterial species in 10 phyla.

### Genomic prediction of metabiotic polysaccharide degradation potential

We first predicted CAZyme gene clusters (CGCs) for each MAG using run_dbCAN v5.0.^54,55^ We retained only CGCs that contained at least one digestive CAZyme (glycoside hydrolase, polysaccharide lyase, or carbohydrate esterase). Because complex polysaccharides are typically too large to cross bacterial cell walls, metabiotic degradation requires that at least one enzymatic step occurs extracellularly or in the periplasm^56^, where breakdown products can diffuse to neighboring cells. We therefore searched for secretion signals by predicting signal peptides in digestive CAZyme proteins using SignalP v6.0^57^ and further restricted the dataset to CGCs in which at least one digestive CAZyme contained a signal peptide.

For each MAG, we counted digestive CGCs with signal peptides and summarized these counts at the family level. Taxonomic assignments followed the Genome Taxonomy Database (GTDB) R226^58^, except for genera traditionally assigned to Prevotellaceae, which GTDB places within Bacteroidaceae. Gut-associated members of Prevotellaceae and Bacteroidaceae in our dataset form reciprocally monophyletic clades^59^, and both family names remain valid according to the List of Prokaryotic names with Standing in Nomenclature as of February 2026^60^; therefore, we treated them as separate families.

To summarize the predicted polysaccharide substrates of CGCs in Bacteroidaceae MAGs, we considered only CGCs that contained a *susC* homolog in addition to the criteria described above. The *susC* gene encodes a TonB-dependent outer membrane porin and is a hallmark of polysaccharide utilization loci in this family^61,62^. SusC homologs were identified using dbCAN annotations of non-CAZyme proteins against the Transporter Classification Database^63^, selecting proteins assigned to TCDB family 1.B.14 that were located within CGCs.

The phylogenetic trees used for data visualization in Figure 6A and 6B are subsets of the bac120 phylogenomic tree from GTDB R226 (https://data.ace.uq.edu.au/public/gtdb/data/releases/release226/226.0/bac120_r226.tree). All plots were generated in R with packages *ggplot, ggtree* (https://doi.org/10.1111/2041-210X.12628), and *treeio*^64^.

### Comparing relative abundance of Bacteroidota families during quinolone exposure

We calculated the median relative abundance of Bacteroidota families per patient, considering only samples from the day after the start of quinolone exposure and up until 30 days after transplantation. Then, we performed a Friedman test to assess overall differences in median abundances across families. For pairwise comparisons between families, we used the paired Wilcoxon signed-rank test, treating each patient as a matched observation across families. Multiple testing correction was applied to the Wilcoxon p-values using the false discovery rate method.

### Predictive modeling of Pseudomonadota BSI with microbiome features

We trained logistic regression classifiers to predict whether a patient had a Pseudomonadota BSI while neutropenic using different microbiome relative abundance features based on 16S sequence data. We compared the predictive performance of models with five different relative abundance feature sets: (i) Log_10_(median Pseudomonadota per patient), (ii) Log_10_(median Bacteroidaceae per patient), (iii) Log_10_(median Tannerellaceae per patient), (iv) Log_10_(median Pseudomonadota per patient) * Log_10_(median Bacteroidaceae per patient), and (v) Log_10_(median Pseudomonadota per patient) * Log_10_(median Tannerellaceae per patient). In all cases, we added a pseudocount of 1×10^-5^ to the median relative abundances to prevent Log_10_(0). We only used samples collected at least one day after the start of quinolone prophylaxis, and up to either the day of engraftment or the day before a positive blood culture of a Pseudomonadota pathogen.

We evaluated model performance using stratified 5-fold cross-validation, ensuring similar proportions of infected and non-infected patients in each fold. In each iteration, models were trained on four folds and tested on the held-out fold, so that every patient contributed to testing exactly once. For test patients, predictions were generated longitudinally using progressively larger subsets of their samples, ordered by collection day relative to transplantation. Specifically, median relative abundances were recalculated using samples collected up to each time point (e.g., up to day −3, day −2,…, day 5), yielding a series of time-dependent predictions for each patient. Predictive performance at each time point was quantified as the area under the receiver operating characteristic curve (ROC-AUC) using the R package *pROC*^65^.

### Imaging and analysis

For experiments involving microfluidics, confocal microscopy was utilized. After the experiment was completed, the devices were left to reoxygenate for 1 hour prior to imaging. Each experiment was then imaged using a Zeiss 880 CLSM with a 40x NA1.4 water objective. 488 and 594 lasers were used to excite the mGFP and mRFP respectively. Representative view fields were chosen at random and imaged with a Z step determined optimal for the objective. These images were then exported for analysis using biofilmQ or visualization using Paraview software.

For dislodged biofilms stained with propidium iodide, samples were taken from each fraction and allowed to reoxygenate for 1 hour prior to imaging 10ul from each fraction was taken and then spotted onto a glass microscope slide and capped with a glass cover slip. Fractions were then imaged at 40x using a Zeiss Axioobserver Z1 with a 40x NA 1.2 air objective. Pa was excited at 488, Bf was excited at 590 and PI was excited at 535nm using a colibri 7. Images were collected with a Hamamatsu camera.

### Quantification and statistics

All image analysis was done using the BiofilmQ image analysis framework^66^. This framework segments fluorescent biomass into cubes with the length of a single cell (2.06um). Analysis can then be performed on these pseudo cells to obtain population level information about the biofilm community. Images are first denoised and then a representative image is thresholded by eye to determine what will be considered biomass. Each channel is then measured by total biomass, height, surface coverage, and other spatial parameters. In gut mimic experiments the “local density” metric was used which measures the % of like biomass within a 20um radius of each individual pseudocell. Data is then exported and figures generated in MATLAB. For pairwise comparisons between data, Nonparametric comparison tests with Bonferroni correction were used. Stars equate a p value of *<0.05, **<0.01, ***<0.001.

**Table S1.** Bacterial strains used in this study.

| Species | Strain | Markers | Reference |
| --- | --- | --- | --- |
| *P. aeruginosa* | PA01 | Chromosomal mGFP | 52 |
| *E. coli* | AD24 | - | 27 |
| *K. pneumoniae* | NCTC 8172 | - | ATCC |
| *B. fragilis* | NCTC 9343 | Chromosomal mRFP | 68 |
| *B. ovatus* | NCTC 11153 | - | ATCC |
| *P. vulgatus* | NCTC 11154 | - | ATCC |
| *B. thetaiotaomicron* | VPI 5482 | - | 68 |

## Acknowledgements

We would like to thank the Sonnenberg lab for their fluorescently labeled *Bacteroides* strains. Research reported in this publication was supported by the National Institutes of Health under project numbers 1R01AI196346-01 and 5P01AI179406-02

## Author Contributions

Conceptualization, B.R.W. and J.B.X; Mouse experiments, S.S, D.K, P.B; Patient data analysis, J.B.X. and I.S; Metagenomic analysis, C.J.P.D; In vitro experiments; B.R.W. and D.K.; Microfluidics and confocal imaging, B.R.W.; Image analysis, B.R.W.; Writing, B.R.W., J.B.X, C.J.P.D. and I.S.; Supervision and Funding, J.B.X.

## Data availability

All processed data used to create the figures of this manuscript are available within the article’s supplementary materials. Raw imaging data are available upon request. Representative processed images and quantification outputs are provided in the supplementary materials. All code for recreating necessary figures can be accessed upon request

