## Supplemental Data for "Metabiosis underlies a microbiota permissive to Pseudomonadota and increases the risk of gut-borne bloodstream infection"

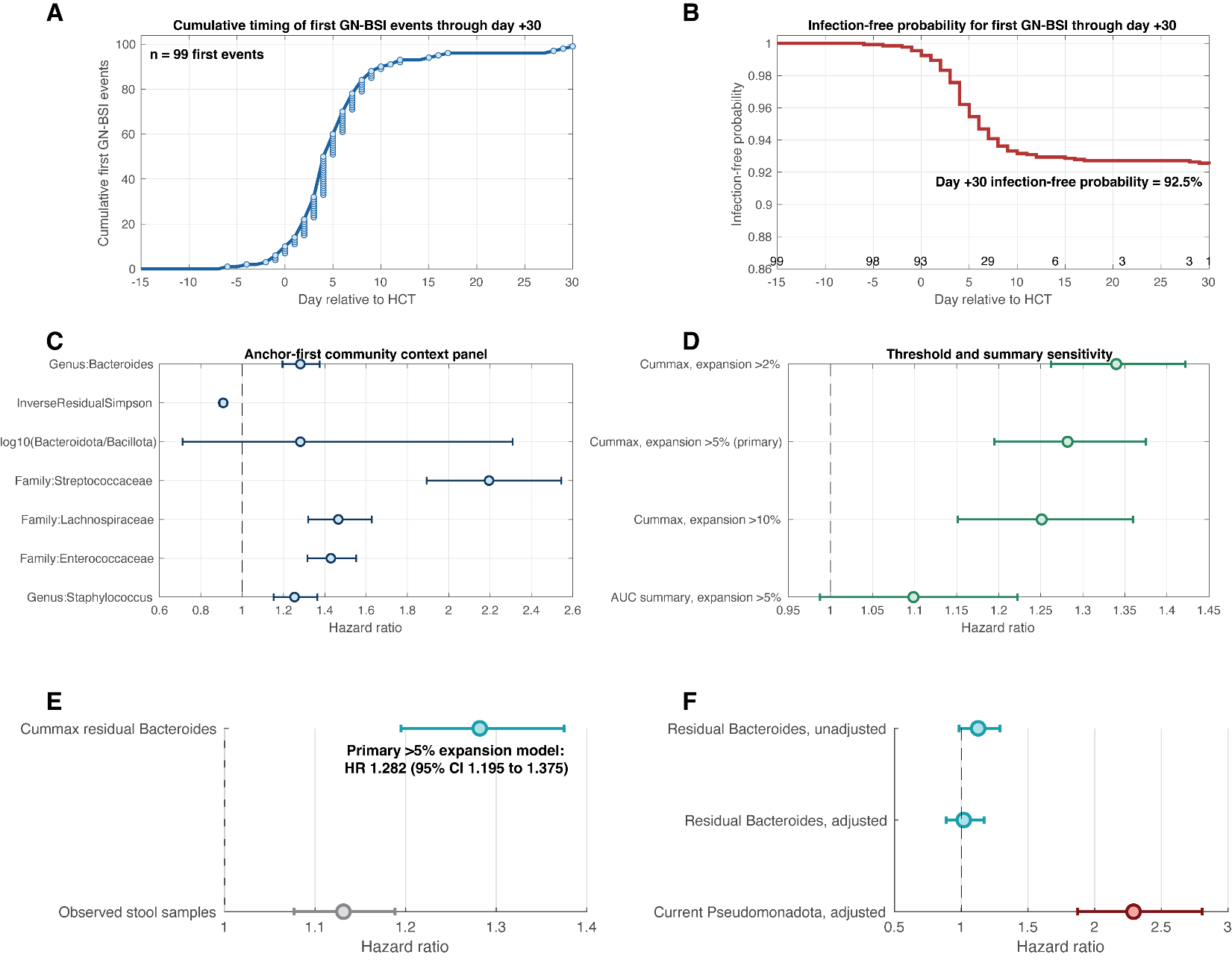

**Figure S1.**

(**A)** Cumulative timing of first Gram-negative bloodstream infection events through day +30. **(B)** Kaplan-Meier panel summarizes infection-free probability through day +30 relative to the transplant day. **(C) Anchor-first community context panel.** Features were fit one at a time in the same start-stop expansion framework, using the exact inferential denominator and interval structure of the primary expansion model. This analysis shows that Bacteroides is part of a broader ecological configuration rather than an isolated variable. (**D) The positive association between residual-*Bacteroides* and later expansion is preserved** across >2%, >5%, and >10% expansion thresholds and remains directionally positive when exposure is summarized by cumulative AUC rather than cummax. The analysis is therefore robust to the operational definition of expansion and to the exact summary statistic used for residual *Bacteroides*. (**E) Peak residual *Bacteroides* is associated with later Pseudomonadota expansion.** Higher peak residual Bacteroides is associated with later ecological takeover after adjustment for cumulative observed sample count. (**F) The residual association between *Bacteroides* and Pseudomonadota attenuates after accounting for the current Pseudomonadota burden.** The pattern is consistent with a pathway in which *Bacteroides* marks a community state that precedes Pseudomonadota expansion, which is then more proximal to GN-BSI.

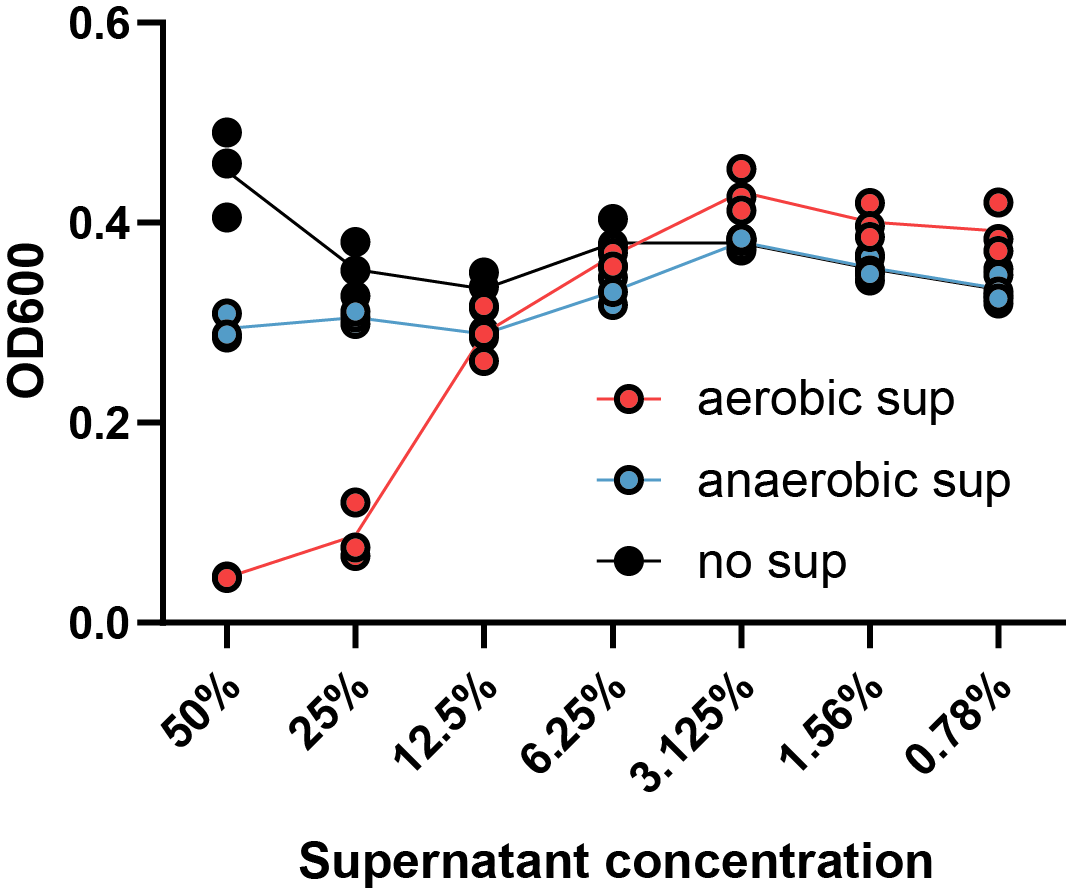

**Figure S2: *P. aeruginosa* supernatant effect on *B. fragilis* growth**

*P. aeruginosa* was cultured in rich media in aerobic and anaerobic conditions before being double filter sterilized and diluted with fresh media before overnight *B. fragilis* culture.

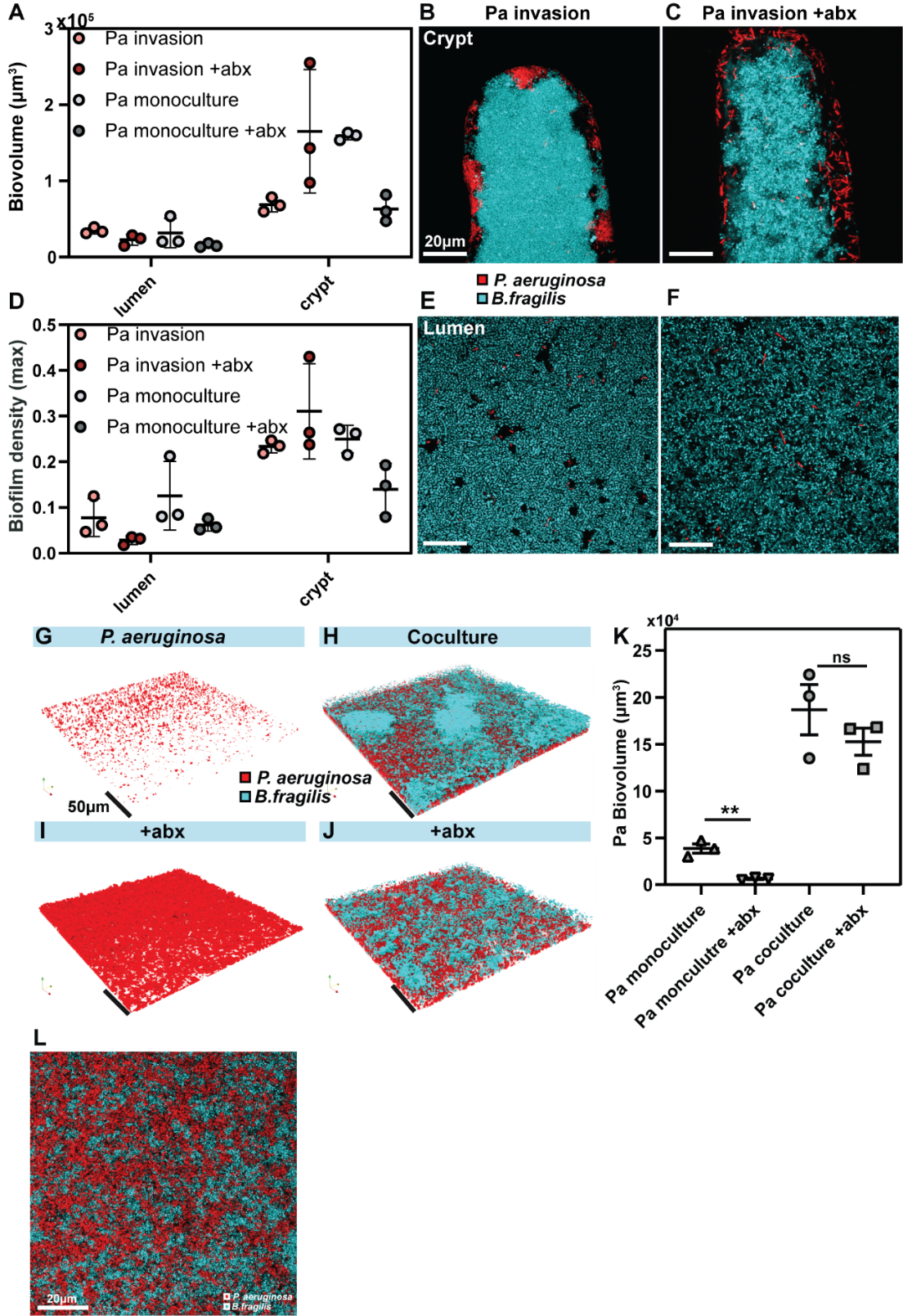

**Figure S3: Crypt invasion and biofilm growth and survival in standard microfluidic culture**

*Bacteroides* biofilms cultured for 48 hours before invasion by *Pseudomonas* and subsequent ciprofloxacin exposure (B/C, E/F). Quantification of biovolume and density of invading *Pseudomonas* biofilm (A,D).3D renderings of biofilms before and after ciprofloxacin treatment in standard microfluidic culture (G-J). Quantification of Pa biovolume before and after ciprofloxacin treatment N=3, Welch’s test **P<.01(K). Representative image showing well mixed nature of Pseudomonadota/*Bacteroides* biofilms.

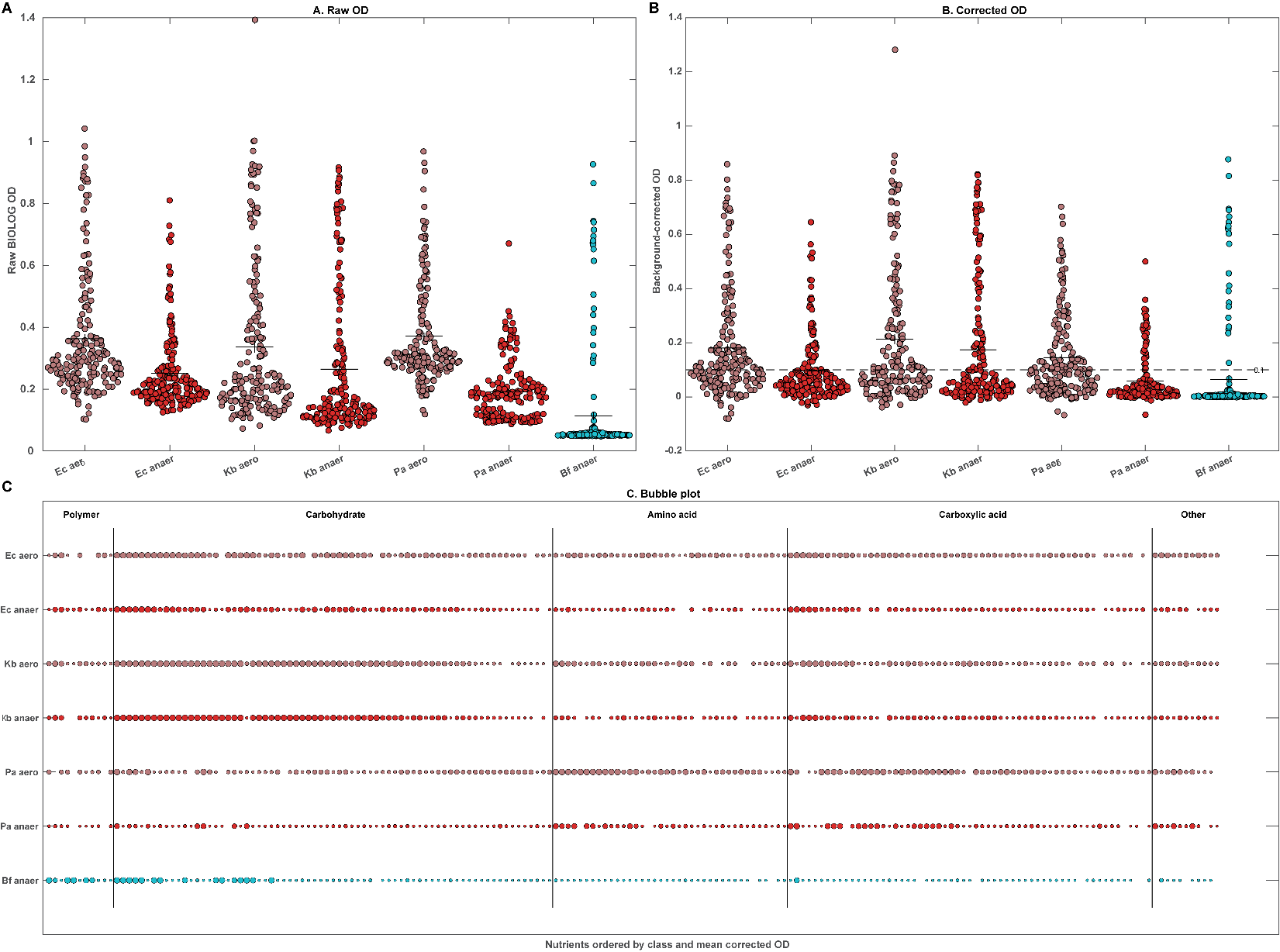

**Supplementary Figure 4. Assay-wide 24 h BIOLOG PM1 and PM2A distributions and full corrected nutrient map underlying Figure 4.**

**(A) Raw OD distributions.** Swarm plots show the distribution of raw BIOLOG OD values across all wells for each strain-condition combination. Horizontal black lines mark the mean and the associated standard error of the mean. These assay-wide distributions provide the uncorrected starting point for the downstream summaries. **(B) Background-corrected OD distributions.** The same well-level distributions are shown after plate-specific background correction. The dashed horizontal line marks the 0.1 utilization threshold used in the Figure 4 breadth and diversity summaries. This panel makes explicit the correction and thresholding framework that underlies the main-text analysis. **(C) Full corrected nutrient map.** All non-control wells are shown after background correction and truncation of negative values to 0. Nutrients are ordered first by broad class and then by mean corrected OD across all strain-condition profiles. Bubble size reflects corrected signal magnitude, and colors follow the manuscript palette. This whole-plate view provides the broader assay context from which the selected main-text fingerprint in Fig. 4A was extracted; full processing details are described in Methods. Canonical methods source: methods.md.

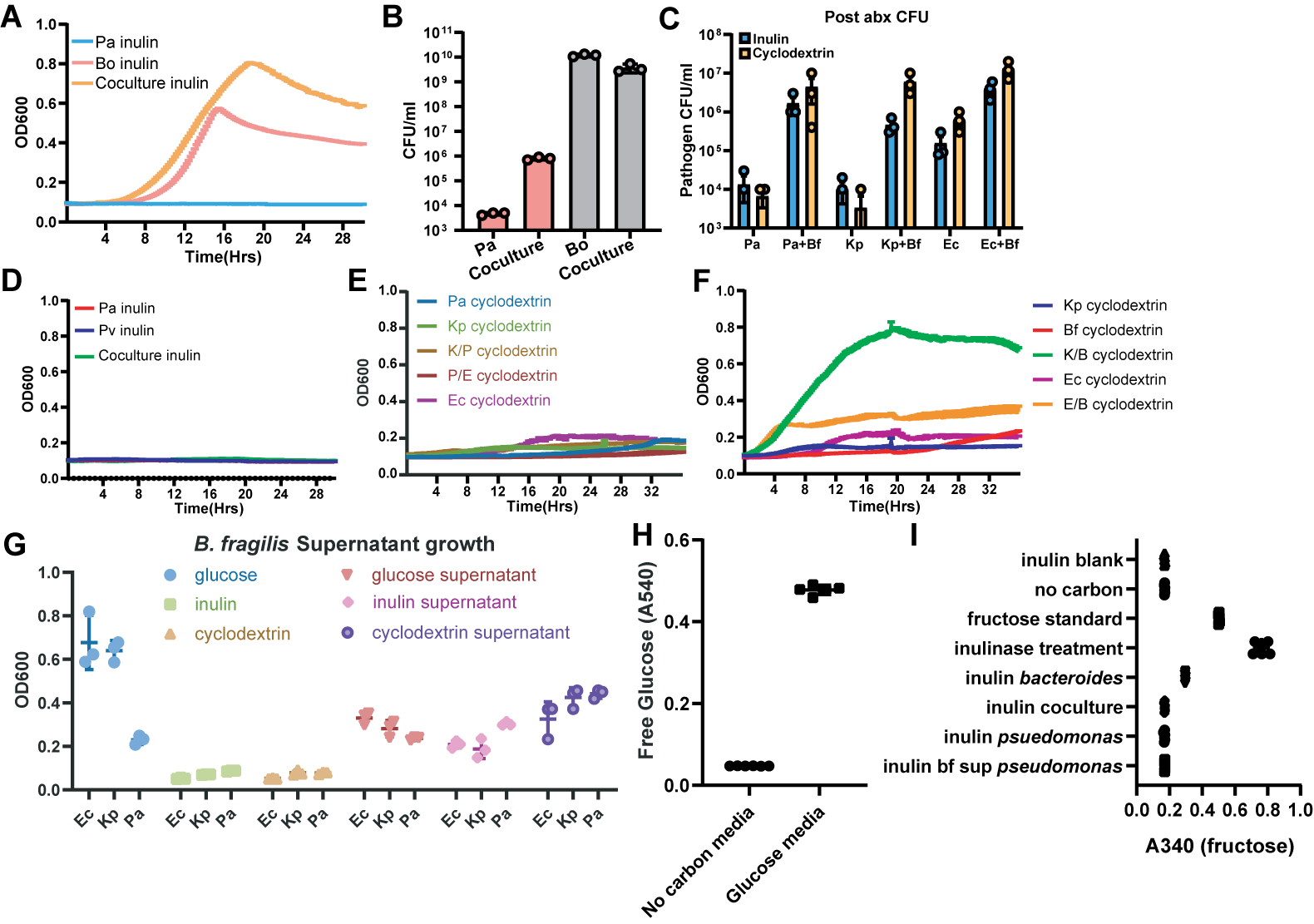

**Figure S5: Metabiotic potential of *Bacteroides* is generalizable across carbon source and species**

**(A) *B. ovatus* can potentiate *P. aeruginosa* on inulin.** Growth curves showing similar kinetics to cyclodextrin. **(B)** CFU counts show same increase in Pa population. **(C) Coculture in cyclodextrin yields antibiotic tolerance across Pseudomonadota.** Each culture was exposed to supra-MIC ciprofloxacin and plated for CFU. **(D/E) Coculture fails to thrive when potentiator is not capable of polysaccharide breakdown.** This is also true of each Pseudomonadota combination. **(F).** Increased speed of growth is seen in coculture across Pseudomonadota. **(G)** cell free *Bacteroides* supernatant is capable of sustaining growth of Pseudomonadota. **(H/I)** controls for free glucose assay and identical experiment to figure 5 done with inulin and free fructose measurement.
